# LC-MS/MS analysis of lesional and normally looking psoriatic skin reveals significant changes in protein metabolism and RNA processing

**DOI:** 10.1101/2020.10.07.329540

**Authors:** V.V. Sobolev, R.H. Ziganshin, A.V. Mezentsev, A.G. Soboleva, M. Denieva, I.M. Korsunskaya, O.A. Svitich

## Abstract

**Background:** Plaque psoriasis is a chronic autoimmune disorder characterized by the development of red scaly plaques. To date psoriasis lesional skin transcriptome has been extensively studied, whereas only few proteomic studies of psoriatic skin are available.

**Aim:** The aim of this study was to compare protein expression patterns of lesional and normally looking skin of psoriasis patients with skin of the healthy volunteers, reveal differentially expressed proteins and identify changes in cell metabolism caused by the disease.

**Methods:** Skin samples of normally looking and lesional skin donated by psoriasis patients (n = 5) and samples of healthy skin donated by volunteers (n = 5) were analyzed by liquid chromatography-tandem mass spectrometry (LC-MS/MS). After protein identification and data processing, the set of differentially expressed proteins was subjected to protein ontology analysis to characterize changes in biological processes, cell components and molecular functions in the patients’ skin compared to skin of the healthy volunteers.

**Results:** The performed analysis identified 405 and 59 differentially expressed proteins in lesional and normally looking psoriatic skin compared to healthy control. We discovered decreased expression of KNG1, APOE, HRG, THBS1 and PLG in normally looking skin of the patients. Presumably, these changes were needed to protect the epidermis from spontaneous activation of kallikrein-kinin system and delay the following development of inflammatory response. In lesional skin, we identified several large groups of proteins with coordinated expression. Mainly, these proteins were involved in different aspects of protein and RNA metabolism, namely ATP synthesis and consumption; intracellular trafficking of membrane-bound vesicles, pre-RNA processing, translation, chaperoning and degradation in proteasomes/immunoproteasomes.

**Conclusion:** Our findings explain the molecular basis of metabolic changes caused by disease in skin lesions, such as faster cell turnover and higher metabolic rate. They also indicate on downregulation of kallikrein-kinin system in normally looking skin of the patients that would be needed to delay exacerbation of the disease. Data are available via ProteomeXchange with identifier PXD021673.

## INTRODUCTION

Psoriasis is a chronic inflammatory skin disorder that affects ∼2%–3% of human population, primarily in the industrialized countries. The most common form of psoriasis is *psoriasis vulgaris* or plaque psoriasis, which is characterized by scaly red skin lesions often located on the scalp, trunk and the extensor surfaces of the elbows and knees. Histology of psoriasis is characterized by incomplete degradation of cell nuclei and intracellular organelles in upper layers of the epidermis, thickening the epidermis due to hyperproliferation of the epidermal keratinocytes, hypogranulosis and massive infiltration of the skin by immune cells, as well as epidermal and vascular remodeling. Although the etiology and pathogenesis of psoriasis remain unclear, nowadays, psoriasis is regarded as an autoimmune disorder, of which the precise mechanism has to be clarified [1, 2].

The inflammatory response in psoriasis is driven by pro-inflammatory cytokines, primarily, interleukin-17, interleukin-23, tumor necrosis factor (TNF) and interferon-γ [3]. For better understanding of the molecular mechanism underlying this process, mRNA profiles from the skin affected by the disease and skin of the healthy volunteers have been compared to each other using microarray and RNA-seq techniques [4-7]. These and other studies have discovered several thousand genes differentially expressed in lesional skin and highlighted the cellular pathways involved in the pathogenesis of psoriasis. They also suggested new treatment options and drug targets [4-7]. In the same time, the changes in mRNA expression often did not correlated with the changes at the posttranscriptional level [8, 9] because the abundance of a protein in proteome, its location and functionality depends on the series of posttranslational events, such as post-translational modification, interactions with binding partners, refolding, proteolytic activation and degradation.

Compared with few thousands of differentially expressed genes (DEGs) revealed in the genomic studies [4-7], the authors of the first proteomic papers that had to rely on two-dimensional electrophoresis achieved very modest results and identified only several dozens of differentially expressed proteins (DEPs) [10-12]. Primarily, it happened due to limited detection range of the used staining method, low reproducibility of the experiments and difficulties in separation of hydrophobic proteins. In contrast, the more advanced LC-MS/MS technique, which is rapidly developing in our time, allowed a detection of more than 1000 DEPs in the same kind of samples [13]. Using this technique in our study we compared the global protein expression in the skin of psoriasis patients and healthy volunteers. Based on the results of performed analysis, we revealed coordinated changes in intracellular signaling and protein and RNA metabolism caused by the disease.

## EXPERIMENTAL

### Ethics statement

All samples were obtained with informed written consent from healthy volunteers and psoriasis patients in accordance with Declaration of Helsinki principles. All protocols were approved by an institutional review board (I.I. Mechnikov Institute of Vaccines and Sera, Moscow, Russia).

### Collection of the skin samples

Skin biopsies were obtained from 5 psoriasis patients: 3 males and 2 females between the ages of 30 and 68 years (mean age 49.2 years) and equal number of healthy volunteers: : 3 males and 2 females between the ages of 39 and 79 years (mean age 61.6 years). The additional details on participants of this study are represented in Table 1. The patients participated in our study discontinued systemic therapies for 2 weeks prior to biopsy collection (1 week in case of topical treatment). Each patient donated two 4 mm punch biopsies of lesional and normally looking (asymptomatic) skin following a local anesthesia. Biopsies of normally looking patients’ skin were taken at least 6 cm away from the nearest skin lesion. The collected biopsies were flash frozen in liquid nitrogen and stored at -85 °C until processing.

**Table 1.** Clinical characteristics of psoriasis patients (n = 5) and volunteers (n = 5) participated in the study.

| ID | Sex | Age | Primary diagnosis | Comorbidities | Medical history |
| --- | --- | --- | --- | --- | --- |
| *Patients* |  |  |  |  |  |
| 1 | male | 68 | plaque psoriasis | psoriatic arthritis | stage 2 hypertension, chronic bronchitis, hyperuricemia, obesity 1st degree |
| 2 | male | 30 | plaque psoriasis |  |  |
| 3 | male | 40 | plaque psoriasis |  | stage 1 arterial hypertension, hyperuricemia, obesity |
| 4 | female | 48 | plaque psoriasis |  | stage 2 hypertension |
| 5 | female | 60 | plaque psoriasis |  | stage 2 hypertension |
| *Volunteers* |  |  |  |  |  |
| 6 | female | 60 |  |  | incisional ventral hernia |
| 7 | female | 79 |  |  | arthrosclerosis, abdominal aortic aneurysm, hypertension, dyslipidemia, chronic pyelonephritis |
| 8 | female | 39 |  |  | white line hernia |
| 9 | male | 77 |  |  | arthrosclerosis |
| 10 | male | 53 |  |  | phlebeurysm |

### Preparation of the samples for analysis

Each sample was washed twice with 0.5 mL phosphate-buffered saline. The samples were homogenized by mechanical disruption in liquid nitrogen. Reduction, alkylation and digestion of the proteins were performed as previously described [14] with minor changes. Briefly, sodium deoxycholate (SDC) lysis, reduction and alkylation buffer pH 8.5, which contained 100 mM TRIS, 1% (w/v) SDC, 10 mM TCEP and 40 mM 2-chloroacetamide was added to the homogenized samples. The samples were sonicated and boiled for 10 min. Then, protein concentration was determined by Bradford assay and the equal volumes of 1% trypsin solution (w/v) prepared in 100 mM TRIS pH 8.5 were added.

After overnight digestion at 37°C, peptides were acidified with 1% trifluoroacetic acid (TFA). The samples (2 x 20 μg) were loaded on 14-gauge StageTips containing 2 layers of SDB-RPS discs. Respectively, 2 tips per a sample were used. The tips were consequently washed with equal volumes of ethyl acetate, 100 μl of 1% TFA prepared in ethyl acetate and 100 μl of 0.2% TFA. After each washing, the excess of liquid was removed by centrifugation (300 g; 1.5 min.) Then, the peptides were eluted with 60 μl of 5% ammonium hydroxide prepared in 80% acetonitrile. The eluates were vacuum-dried and stored at -80°C. Prior the experiment, the vacuum-dried samples were dissolved in 2% acetonitrile/0.1% TFA buffer) and sonicated for 2 min.

### LC/MS-MS analysis

The reverse-phase chromatography was performed on Ultimate 3000 Nano LC System (Thermo Fisher Scientific) coupled to the Q Exactive Plus benchtop Orbitrap mass spectrometer (Thermo Fisher Scientific) using a chip-based nanoelectrospray source (Thermo Fisher Scientific). The columns were packed in the lab as previously described [15]. Samples prepared in the loading buffer (0.1% TFA and 2% acetonitrile in water) were loaded on Inertsil ODS3 (GLSciences, USA) trap column (0.1 x 20 mm, 3 μm), at 10 μl/min and separated on Reprosil PUR C18AQ (Dr. Maisch, Germany) fused-silica column (0.1 x 500 mm, 1,9 μm) with linear gradient of 3-35% buffer B (0.1% formic acid, 80% acetonitrile in water) for 55 min; 35-55% B for 5 min, and 55-100% B for 1 min at flow rate of 440 nl/min. Prior to the next sample injection, the column was washed with buffer B for 5 min and reequilibrated with buffer A (0.1% formic acid, and 3% acetonitrile in water) for 5 min.

Peptides were analyzed on the mass spectrometer with one full scan (350–2000 *m*/*z, R* = 70,000 at 200 *m*/*z*) at a target of 3 × 10^6^ ions and max ion fill time 50 ms, followed by up to 10 data-dependent MS/MS scans with higher-energy collisional dissociation (HCD) (target 1 × 10^5^ ions, max ion fill time 45 ms, isolation window 1.4 m/z, normalized collision energy (NCE) 27%), detected in the Orbitrap (*R* = 17,500 at fixed first mass 100 *m*/*z*). Other settings: charge exclusion - unassigned, 1, more than 6; peptide match – preferred; exclude isotopes – on; dynamic exclusion 40 s was enabled.

### Analysis of LC-MS data

Label-free protein quantification was performed using MaxQuant software version 1.5.6.5 [16], and a common contaminants database by the Andromeda search engine [17] with cysteine carbamidomethylation as a fixed modification. Oxidation of methionine and protein N-terminal acetylation were used as variable modifications. Peak lists were searched against the human protein sequences extracted from the Uniprot (28.06.19) database. The false discovery rate (FDR_ was set to 0.01 for both proteins and peptides with a minimum length of seven amino acids. Peptide identification was performed with an allowed initial precursor mass deviation up to 20 ppm and an allowed fragment mass deviation of 20 ppm. Downstream bioinformatics analysis was performed using Perseus [18] (versions 1.5.5.1). Protein groups only identified by site, only from peptides identified also in the reverse database, or belonging to the common contaminants database were excluded from the analyses. For Student’s t-test, missing values were imputed with a width of 0.3 and a downshift of 1.8 over the total matrix. Two sample tests were performed in Perseus with s0 set to 0. Label free quantification was performed using a minimum ratio count of 1 [19]. The protein levels were assessed by the iBAQ (intensity-based absolute quantification) method using MaxQuant software [20]. To determine the relative abundance of identified proteins in the samples (riBAQ), we divided the obtained iBAQ values by the sum of all iBAQ values, and expressed this ratio as a percentage [21]. The proteins were considered as differentially expressed if they exhibited a fold-change of at least 1.5 and FDR was less than 0.05. The results were analyzed using Venn diagrams and heat maps.

Protein ontology analysis of the differentially expressed proteins (PO) was performed on gene ontology terms to catalog the molecular functions, cellular components and biological processes using DAVID Bioinformatics resources, 6.7 (Frederick National Laboratory for Cancer Research, USA).

### Statistical analysis

Data are shown as mean ± SE and were analyzed with a two-sided, unpaired Student’s t-test. Differences were considered statistically significant when p < 0.05. The mass spectrometry proteomics data were deposited to the ProteomeXchange Consortium via the PRIDE [22] partner repository with the dataset identifier PXD021673.

## RESULTS AND DISCUSSION

### Proteomes of normally looking and lesional skin of psoriasis patients are different from the skin proteome of healthy volunteers

LC/MS-MS analysis of the skin samples taken from psoriasis patients (n = 5) and healthy volunteers (n = 5) identified 899 proteins (Table S1) and 520 proteins were differentially expressed (Table S2). The numbers of identified proteins and DEPs was comparable with other proteomic studies [9, 13]. The distribution of these proteins between the groups is shown as a Venn diagram (Fig. 1). The set of proteins located in the center of the diagram consisted of 16 DEPs that were differentially expressed regardless what two groups of samples we compared. Since the expression of 8 DEPs, namely S100A7, RPL39, NACA, RAB1A, RAB11B, CKMT1A, AHCY and TGM3, progressively raised in normally looking skin of the patients compared to healthy skin and in lesional skin of the patients compared to their normally looking skin, we would consider them as potential disease biomarkers. Presumably, the expression levels of the named DEPs could be used to identify the individuals predisposed to psoriasis and monitor the disease progression in the patients. In contrast, the three peripheral sets of 12, 59 and 82 proteins were differentially expressed when we compared two specific groups of samples. Respectively, any set of these DEPs could be used to describe differences between these two specific groups. For instance, the set of 82 DEPs (Fig. 1) could help to characterize the transition from normally looking to lesional skin. In the other words, it could be used to track the development of psoriatic plaque. In turn, the remaining three sets of 14, 25 and 312 proteins distinguished a specific group of samples from the others. For instance, 312 proteins mentioned above (Fig. 1) were differentially expressed in lesional skin when the samples obtained from lesional skin were compared to the samples obtained from either normally looking skin or skin of the healthy volunteers. However, these proteins were not differentially expressed when samples obtained from normally looking skin of the patients were compared to the samples obtained from skin of the healthy volunteers. Thus, the mentioned 312 DEPs could be used to characterize the specific skin condition, which would be lesional skin.

**Figure 1.**
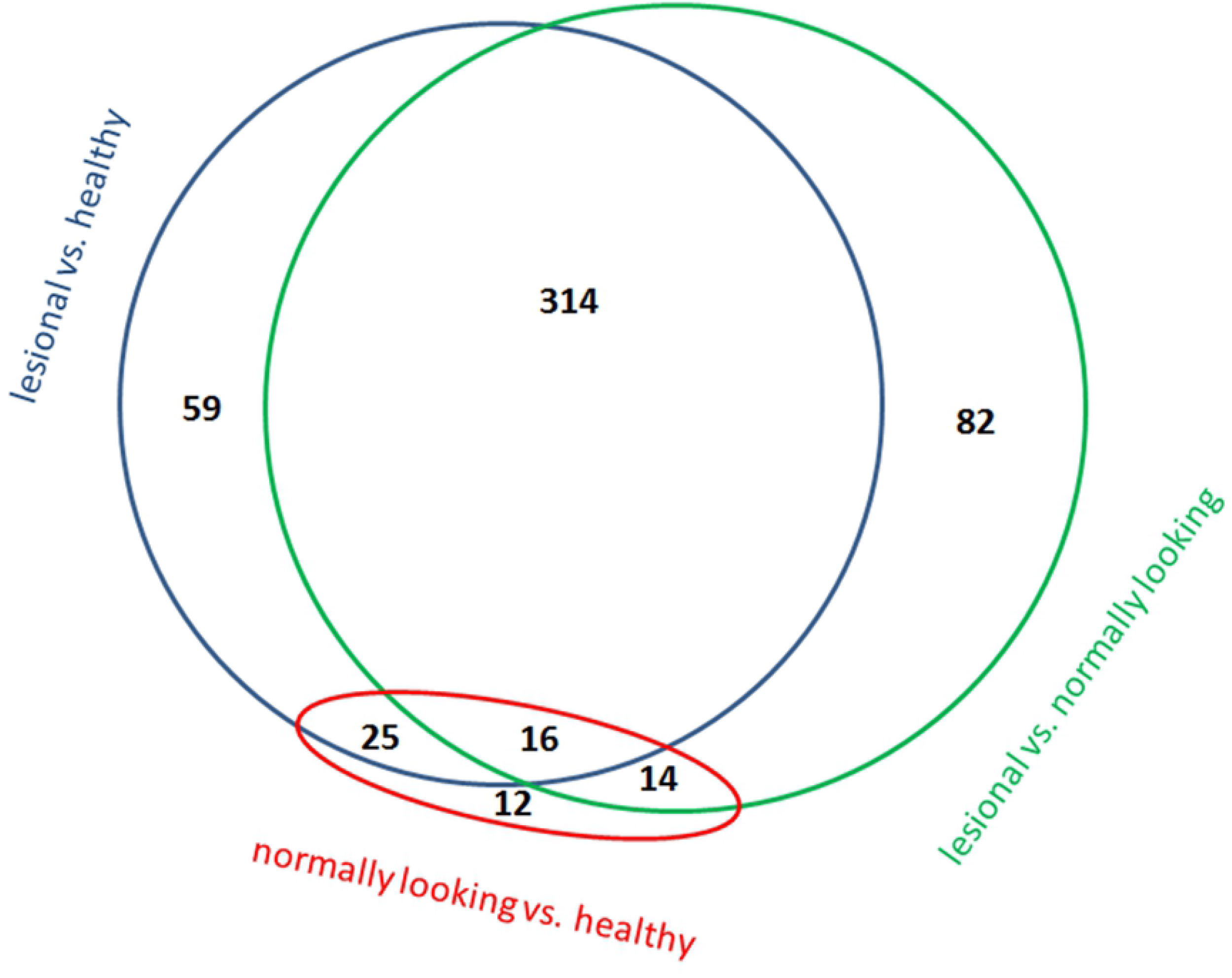
Venn diagram comparing DEPs in skin samples obtained from lesional and normally looking skin of the same psoriasis patients (n = 5) andskin of the healthy volunteers (n = 5) as assessed by LC/MS-MS analysis. The numbers indicated in the diagram are the numbers of DEPs in compared groups of samples (FDR < 0.05, fold change >1.5).

### Proteins differentially expressed in uninvolved skin were overrepresented in extracellular compartments and secretory granules

When normally looking skin of psoriasis patients was compared to the skin of the healthy volunteers, 59 proteins were differentially expressed. PO analysis of biological processes (Table S3) revealed their enrichment in platelet degranulation (P = 0.0076). In turn, PO analysis of cellular components identified 10 terms (Table S4). The highest enrichment was for extracellular exosomes (P = 5.10 x 10^−16^), blood microparticles (P = 1.60 x 10^−14^) and platelet α granule lumen (P = 1.60 x 10^−4^). The largest mentioned group (platelet degranulation) included two others, except HBD that was associated only with “platelet α granule lumen”. This result suggested us that major differences of the patients’ normally looking skin from healthy skin of the volunteers were presumably caused by differential expression of proteins involved in intercellular signaling, and particularly ones that were associated with specialized granules, such as extracellular exosomes. In turn, when the DEPs were analyzed for overrepresentation in terms of molecular functions, all identified terms exhibited FDR > 0.05 (Table S5).

When PO analysis of 26 DEPs with increased expression was performed separately, no significant enrichment in terms of biological processes or molecular functions was seen. In terms of cellular components (Table S4a), we discovered enrichment for extracellular exosomes (P = 6.10 x 10^−4^). In contrast, PO analysis of 33 DEPs with decreased expression (Table S3a) revealed enrichment for two biological processes, namely regulation of blood coagulation (P = 4.70 x 10^−4^) and regulation of coagulation (P = 4.00 x 10^−4^). In both processes, DEPs were the same (KNG1, APOE, HRG, THBS1 and PLG). Taking in account their roles in the cell, we would suggest that their suppression could slow down the activation of kallikrein-kinin system and the recruitment of the immune cells to the sites of inflammation. Analyzing the cellular components (Table S4b), we discovered that DEPs were enriched in extracellular region (P = 6.20 x 10^−3^; FDR = 0.055). In turn, the analysis of molecular functions (Table S4a) revealed that DEPs were enriched in heparin (P = 2.00 x 10^−4^) and glycosaminoglycan (P = 4.50 x 10^−4^) binding, which was not surprise considering the active role glycoproteins in binding of immunocytes and proinflammatory cytokines (e.g. CXCL8/IL8) to vascular endothelial cells. Thus, the threshold of the inflammatory response in normally looking skin of the patients supposed to be higher than in the skin of healthy volunteers. This phenomenon could be a part of an adaptation mechanism aimed to delay the exacerbation of the disease.

### The proteins differentially expressed in lesional skin are responsible for higher metabolic rate accelerating pre-RNA processing, translation, protein biosynthesis and degradation

A comparative analysis of the skin samples obtained from lesional skin of the psoriasis patients and the skin of the healthy volunteers revealed 405 differentially expressed proteins. The expression levels of 331 proteins were significantly increased whereas the expression levels of 74 proteins were significantly decreased. The DEPs with highest expression in lesional skin were S100A7, SERPINB3 and NAMPT (Table 2).

**Table 2.** The most abundant proteins of lesional skin

| **Protein** | **Uniprot ID** | **P-value** | **Fold increase in lesional skin\*** |
| --- | --- | --- | --- |
| S100A7 | P31151 | 1.33E-05 | 80.10 |
| SERPINB3 | P29508 | 4.48E-03 | 67.86 |
| NAMPT | A0A024R718 | 6.39E-05 | 45.08 |
| SCCA1-SCCA2 | Q5K684 | 2.16E-03 | 35.60 |
| TYMP | P19971 | 1.25E-03 | 29.56 |
| A2ML1 | A8K2U0 | 5.96E-05 | 26.10 |
| S100A8 | P05109 | 1.12E-05 | 25.80 |
| KRT16 | P08779 | 3.49E-02 | 25.47 |
| TGM3 | Q08188 | 1.44E-03 | 23.36 |
| FABP5 | Q01469 | 8.79E-06 | 22.17 |
| IDE | A0A3B3ISG5 | 6.99E-06 | 21.35 |
| RPL39 | P62891 | 2.15E-06 | 18.49 |
| S100A2 | P29034 | 3.96E-03 | 17.63 |
| IVL | P07476 | 1.01E-03 | 14.81 |
| AKR1B10 | O60218 | 1.31E-05 | 12.67 |
| GLUL | A8YXX4 | 2.01E-02 | 12.38 |
| LCN2 | P80188-2 | 9.64E-03 | 12.03 |
| S100A9 | P06702 | 8.77E-05 | 11.78 |
| GSDMA | Q96QA5 | 3.08E-03 | 11.21 |
| STAT1 | P42224 | 2.69E-05 | 10.82 |

\* The expression level in skin of the healthy volunteers was equal to 1

Previously, three different groups of researchers compared the skin samples obtained from psoriasis patients and healthy volunteers using 2D gel electrophoresis followed by MS. Piruzian et al. [11] reported of the 10 most abundant proteins in skin lesions. All these proteins were also identified in our study (Table S2). Particularly, we confirmed the elevated expression of KRT16, SERPINB4, SERPINB3, S100A9, S100A7, KRT17 and ENO1. Moreover, the first five of them exhibited the highest expression levels in lesional skin of our patients (Table 2). On the other hand, the expression of KRT14, ENO1 and LGALS7B did not exceed 2 or 3 fold. In this regard, we would like to mention that differences between their and our data could be caused by technical problems with scanner and densitometer. Moreover, the band intensities did not suppose to grow proportionally to protein amounts in the samples if they tended to approach the upper detection limit.

Carlén et al. [10] identified 11 DEPs in lesional psoriatic skin. In their paper, the authors reported of 9 proteins those expression was increased and 2 proteins those expression was decreased, compared to the skin of the healthy volunteers. Unlike us, they did not detect KRT10 in lesional skin that had to be abundant in stratified epithelia (Table S2). In turn, we could not identify KRT15 that was detected in their study. Moreover, we had a problem with interpretation of the terms “SCCA2, low pI” and “SCCA2, high pI” that Carlén et al. used as “protein names”. This problem could be avoided if they provided valid Uniprot IDs. After all, we confirmed the expression of SERPINB5/maspin, KRT17, GSTP1/GST-π, HSPB1/HSP27, ARHGDIA/RhoGDI-1, CALR/calreticulin and SFN/14–3–3 σ, although according to our data, only KRT17 (P=0.040), GSTP1 (P=0.018) and CALR (P=0.004) could be differentially expressed.

In turn, Ryu et al. [12] identified 30 DEPs in skin lesions of psoriasis patients, compared to their normally looking skin. Comparing their data with ours, we could not confirm the differential expression of TF, ATP5B, HSP90B1, PARK7, PDIA3, APCS and SERPINF1. In turn, four other proteins, namely DPYSL2, HBB, PRDX2 and SERPINA1 that Ryu et al. reported as proteins exhibiting an increased expression were suppressed in the skin of our patients. On the other hand, we confirmed the elevated expression of 17 other proteins, including GSTP1 (P = 0.018), which was reported by them as DEP for the first time. In addition, the expression levels of the five remained proteins reported by them (HSPB1, MDH1, SFN, TUBB2C and YWHAZ), changed less than 1.5-fold in our study. In addition, we were not sure whether we and their group used the same criteria to identify DEPs because they did not provide any quantitative estimate of protein expression levels in their paper.

Comparing lesional skin and healthy control, we identified several DEPs controlling the terminal differentiation of epidermal keratinocytes. Particularly, we found that the expression level of the early differentiation marker IVL was increased (14.81 ± 5.36, P = 0.001), whereas the expression level of the late differentiation marker FLG was decreased (0.50 ± 0.16, P = 0.016). We also discovered the bidirectional changes in the expression of cytokeratins. The expression levels of KRT1 and KRT10 were decreased (0.58 ± 0.11, P = 0.011 and 0.56 ± 0.09, P = 0.005, respectively). In this regard, we would like to mention that KRT1 and KRT10 are the binding partners that interact in 1:1 ratio. In the cell, their expression had to be coordinated and changes in their expression levels be comparable to each other. The expression levels of two other cytokeratins KRT16 (25.47 ± 18.69, P = 0.035) and KRT17 (2.85 ± 1.38, P = 0.090) were increased.

Moreover, we identified three other cytokeratins, which were not abundant in healthy skin. In lesional skin, the expression of KRT23, the paralog of KRT14 in the human genome, was increased (2.79 ± 1.01, P=0.002). In contrast, the expression levels of KRT72 and KRT77 were decreased (0.49 ± 0.21, P=0.05 and 0.04 ± 0.03, P=0.003). Although the role of KRT72 and KRT77 in the epidermis remains uncharacterized, their lower expression levels in lesional skin compared to healthy control were already confirmed by the others on both protein and mRNA levels [23, 24].

In addition, we showed an increased expression of psoriasin/S100A7 (80.10 ± 25.67, P=10^−5^), S100A8 (25.80 ± 8.45, P=10^−05^) and S100A9 (11.78 ± 8.47, P=8E-05). According to the others [13, 23, 24], the named proteins were overexpressed in lesional skin. Unfortunately, we did not identify the differential expression of the proliferation marker MKI67 that supposed to be significantly increased and the late differentiation marker LOR that had to be decreased in lesional skin.

The PO analysis showed enrichment of 46 biological processes with FDR < 0.05 (**Table S6**). The highest enrichment was observed in translation elongation and translation (P = 1.60 x 10^−62^ and 1.80 x 10^−48^ respectively), positive regulation of ubiquitin-protein ligase activity, regulation of ubiquitin-protein ligase activity during mitotic cell cycle (P = 3.10 x 10^−10^ and 4.00 x 10^−10^ respectively) and positive regulation of ligase activity (P = 5.50 x 10^−10^). When PO analysis was performed separately on DEPs that exhibited an increased expression (Table S6a), the most enriched biological processes kept the same order.

It is well-known that development of psoriatic plaques is accompanied by a genome-wide shift in expression of several thousand DEGs [9] and rapid adjustment of the regulatory apparatus for translation of the transcribed mRNAs into the proteins. Since the DEPs involved in the translation had different roles, we split them in several subgroups. Primarily, we recognized 23 and 31 proteins of the small and large ribosome subunits. Most of them were underexpressed in the patients’ normally looking skin (Fig. 2a). In contrast, their expression levels were significantly increased in lesional skin of the same patients. Similarly, 10 differentially expressed aminoacyl-tRNA synthetases (Fig. 2b) and 21 factors that controlled initiation, elongation and termination of translation (Fig. 2c) were suppressed in healthy looking skin and elevated in skin lesions, compared to healthy skin.

**Figure 2.**
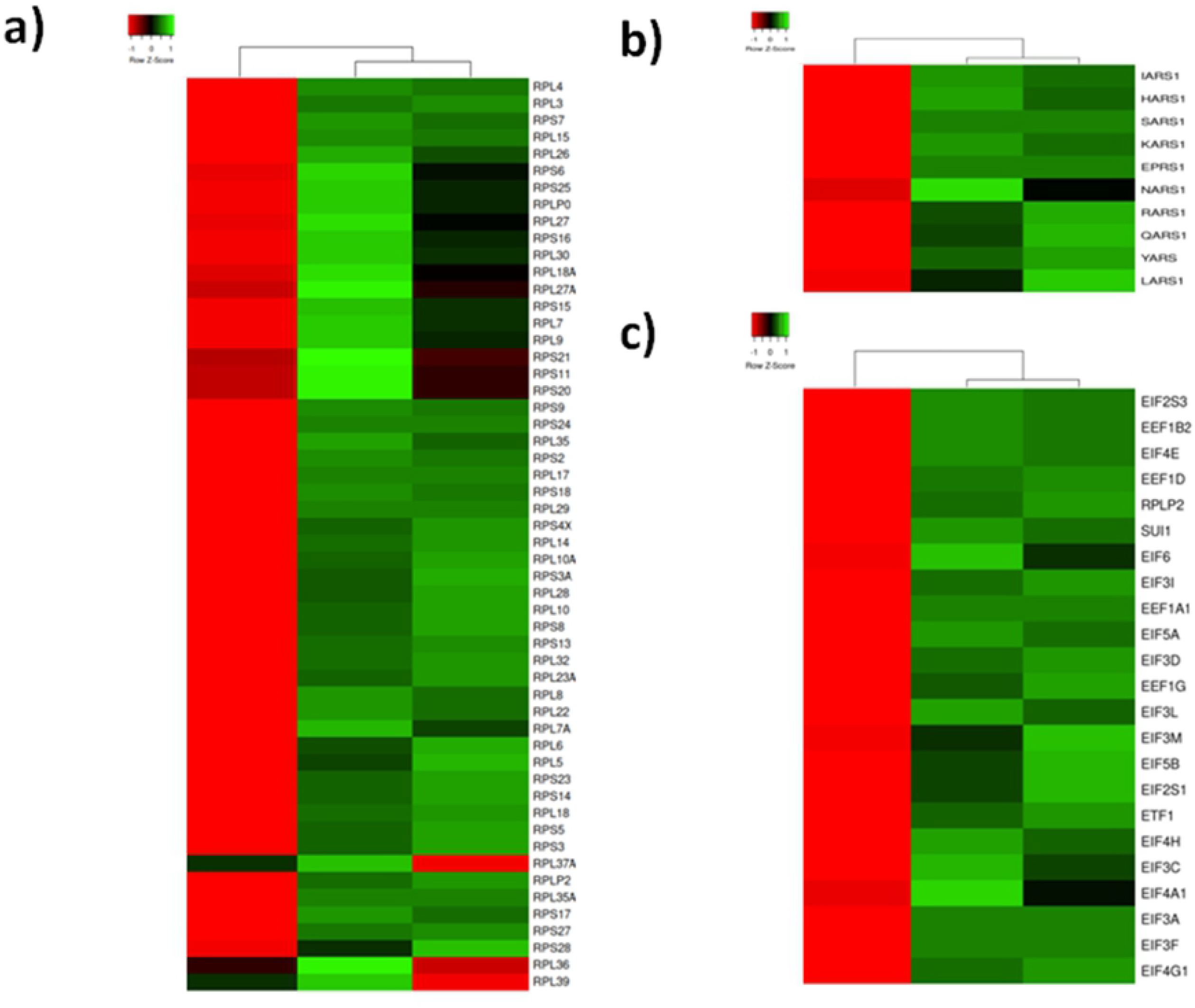
Heat map representing three clusters of DEPs involved in mRNA translation, according to the average quantitative expression levels in two groups of samples. a) Ribosome proteins; b) Aminoacyl-tRNA synthetases; c) Translation initiation elongation and termination factors; From left to right column – the results of comparative analysis for normally looking skin of the patients vs. skin of the healthy volunteers, lesional vs. normally looking skin of the patients, and lesional skin of the patients vs. skin of the healthy volunteers. Red color indicates low expression levels whereas green color indicates high expression levels. The DEPs uniprot ID, official gene symbols of the encoding genes, fold changes and Benjamini corrected P-values are shown in Supplementary Table S2.

These findings were supported by the other researches. A long time ago, Freedberg reported a 7-fold increase of translation in lesional skin [25] and explained it by a delay in degradation of ribosomal proteins. Recently, Swindell et al. discovered an elevated expression of translation-related proteins, primarily the ribosome proteins, in lesional skin [9]. In turn, we noticed that about a half of aminoacyl-tRNA synthetases and multiple translation factors were also differentially expressed. Based on these findings, we would like to hypothesize that major changes in psoriatic proteome were needed to maintain a higher metabolic rate in lesional skin.

Similarly to the other methods, ontology analysis has own limitations and some of its results could be controversial. For instance, about a half of biological processes that we presented in Table S6 were enriched by the same group of DEPs, which was mainly composed of proteasome and immunoproteasome proteins. These biological processes were associated with different aspects of ubiquitin-protein ligase activity and its regulation. Some of these biological processes, like negative and positive regulation of protein ubiquitination (P = 3.10 x 10^−10^ and 8.10 x 10^−10^ respectively) supposed to have the opposite outcomes. To our surprise, the same DEPs, except SKP1, which was associated only with the first one, were enriched in both of them. Taking in account that SKP1 catalyzed the ubiquitination of proteins designated for proteasomal degradation while the other DEPs (proteasome and immunoproteasome subunits) directly participated in the 26S proteasomal degradation pathway, we would consider a positive regulation of protein ubiquitination as the only true outcome and, respectively, disregard negative regulation of protein ubiquitination.

To the notice, the expression pattern of proteasome/immunoproteasome subunits (Fig. 3) looked similar to the pattern of DEPs involved in translation (Fig. 2). Most of them exhibited decreased expression in normally looking skin and increased expression in lesional skin of the patients, compared to the control group. Particularly, the mentioned group of DEPs consisted 6 of 7 α-subunits (PSMA1-6), two β-subunits (PSMB1 and PSMB7) shared by proteasome and immunoproteasome, one β-subunit of immunoproteasome (PSMB8, a substitute of the non-mentioned β-subunit of proteasome PSMB5), 3 subunits exhibiting ATPase activity (PSMC3, PSMC4 and PSMC5), 4 subunits (PSMD2, PSMD6, PSMD11 and PSMD12) that did not exhibit ATPase activity and belonged to the regulatory module of proteasome (19S) and two subunits of E-type (PSME1 and PSME2) that composed the regulatory module of immunoproteasome (11S). Since all of them exhibited increased expression in lesional skin, we suggested a balance shift between 11S and 19S in favor of 11S because both its subunits (PSME1 and PSME2) were increased in skin lesions. Moreover, we would suggest an activation of both: proteasome and immunoproteasome. The former made sense due to sustained skin inflammation is the hallmark of psoriasis. The latter favored our hypothesis that changes in protein expression primarily directed to please a higher metabolic rate in the skin cells affected by the disease. The obvious way to achieve it would be to accelerate the metabolic processes running in the opposite direction (e.g. translation and protein degradation in proteasomes).

**Figure 3.**
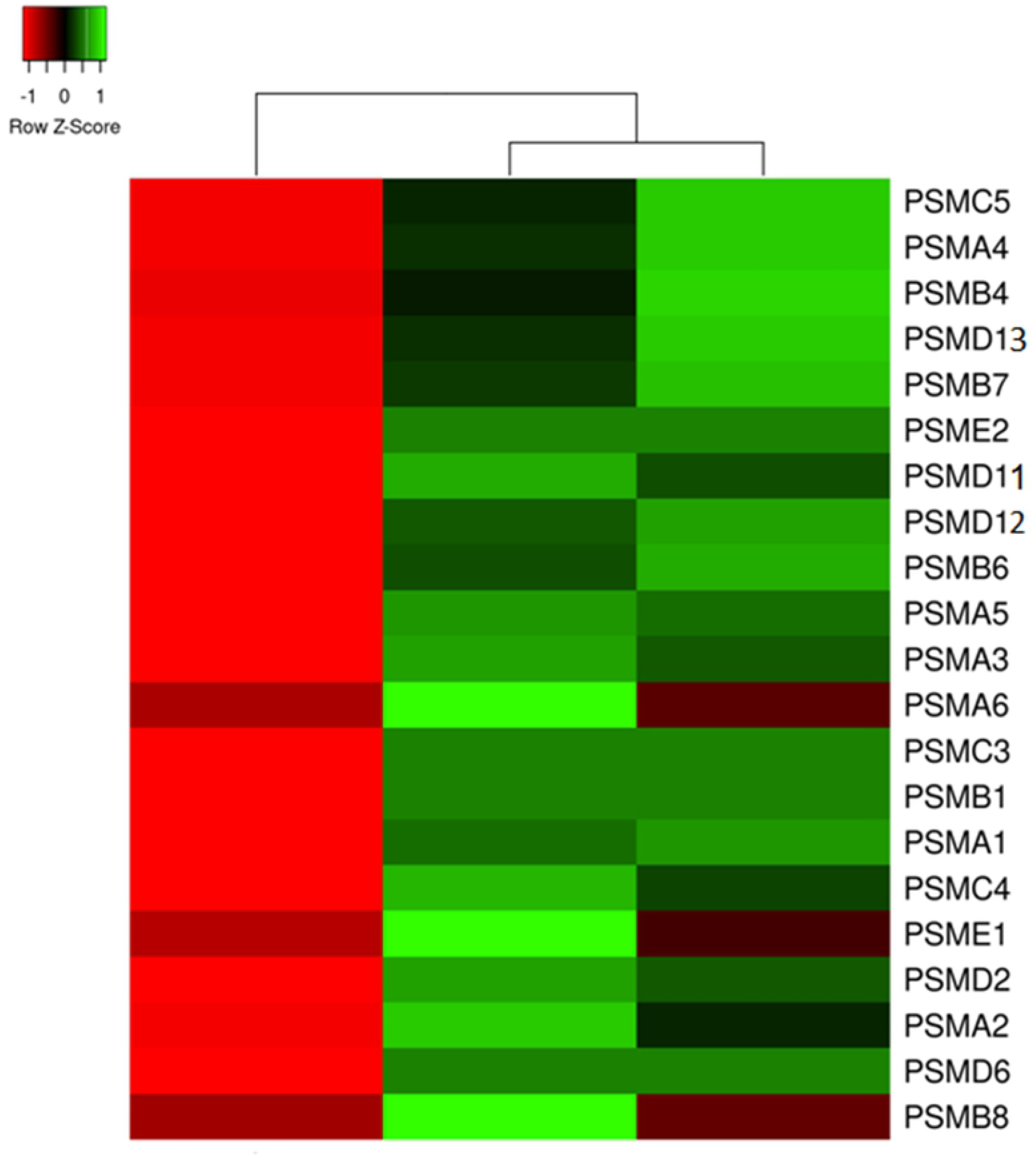
Heat map representation of the differentially expressed subunits of proteasome and immunoproteasome, according to the average quantitative expression levels in two groups of samples. From left to right column – the results of comparative analysis for normally looking skin of the patients vs. skin of the healthy volunteers, lesional vs. normally looking skin of the patients, and lesional skin of the patients vs. skin of the healthy volunteers. Red color indicates low expression levels whereas green color indicates high expression levels. The DEPs uniprot ID, official gene symbols of the encoding genes, fold changes and Benjamini corrected P-values are shown in Supplementary Table S2.

Enrichment analysis applied to the GO cellular component terms (Table S7) showed the highest representation for cytosol (P = 6.32 x 10^−56^), cytosolic ribosome (P = 3.93 x 10^−54^) and cytosolic parts (P = 3.98 x 10^−50^). The terms proteasome complex (P = 3.12 x 10^−12^), membrane-bound vesicle (P = 1.80 x 10^−4^) and melanosome (P = 2.11 x 10^−10^) were also enriched. In turn, the DOPs exhibiting an increased expression were enriched the same terms (Table S6a), except eukaryotic translation initiation factor 3 complex (P = 1.67 x 10^−4^), which was not enriched when all DEPs regardless their expression level were analyzed.

Similarly to the DEPs enriched in translation and proteasomal degradation (see above), most of the DEPs enriched in membrane-bound vesicles exhibited decreased expression in normally looking skin and increased expression in lesional skin of the patients (Fig. 4). Primarily, we would like to mention two adaptin proteins (AP2B1 and AP2M1) that contributed to the AP-2 complex located on the surface of clathrin-coated vesicles. We would also mention CLTC, which was the heavy chain of clathrin. Moreover, we would acknowledge differential expression of several small GTPase that regulate the functioning of the clathrin-coated vesicles. One of them, RAB14 was found in vesicles that carried newly synthesized proteins from the Golgi apparatus to early endosomes. In turn, RAB11B controlled the performance of exo- and endocytosis. RAB5C secured docking and fusion of the vesicles to a target membrane. RAB7A regulated the formation of late endosomes as well as their transition to lysosomes. Since all these enzymes were strongly expressed in lesional skin, it would be reasonable to propose an activation of the protein trafficking toward the primary endosomes and lysosomes. This assumption is in a good agreement with our hypothesis that changes in psoriatic proteome are primarily needed to support a faster metabolic rate in the disease-affected skin because it links together protein biosynthesis and protein degradation.

**Figure 4.**
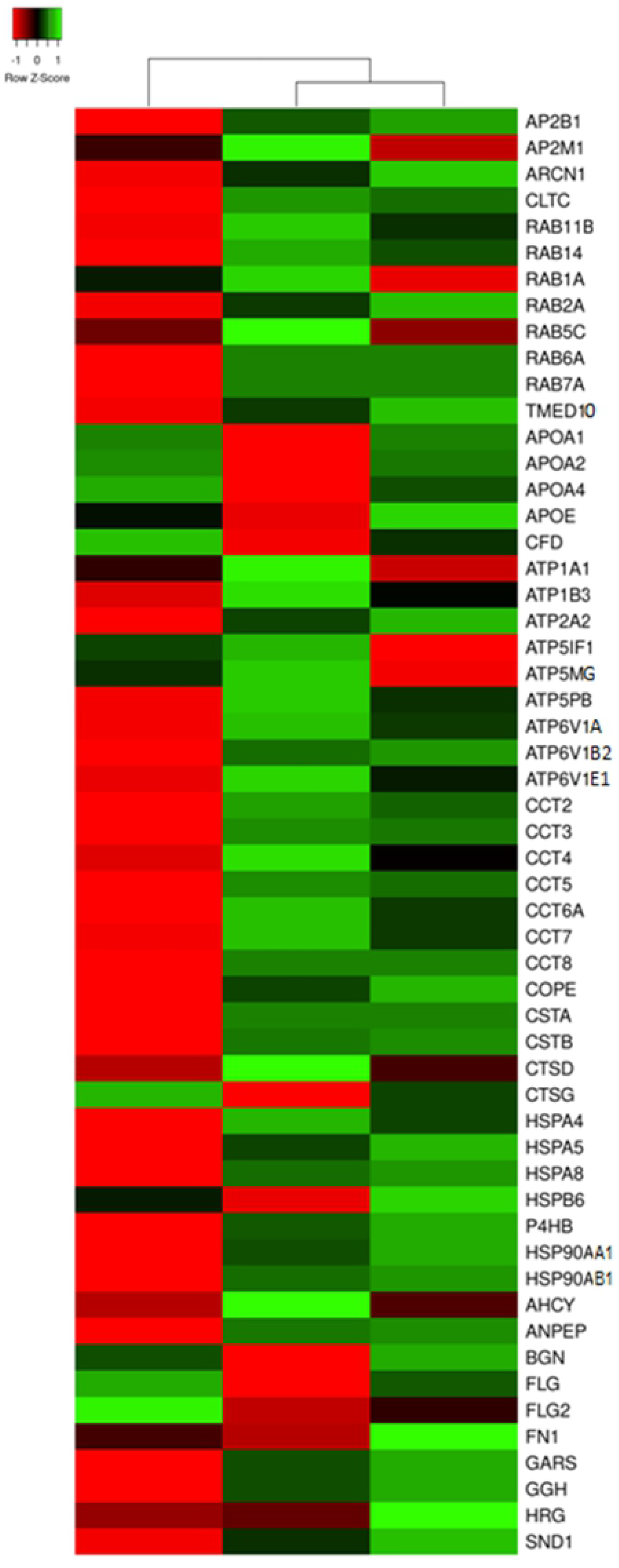
Heat map showing mean expression profiles of the DEPs associated with membrane-bound vesicles, according to the average quantitative expression levels in two groups of samples. From left to right column – the results of comparative analysis for normally looking skin of the patients vs. skin of the healthy volunteers, lesional vs. normally looking skin of the patients, and lesional skin of the patients vs. skin of the healthy volunteers. Red color indicates low expression levels whereas green color indicates high expression levels. The DEPs uniprot ID, official gene symbols of the encoding genes, fold changes and Benjamini corrected P-values are shown in Supplementary Table S2.

Expectedly, we observed significant changes in expression of lysosomal proteins. Most of them exhibited decreased expression in normally looking skin and increased expression in lesional skin of the patients. Three of these proteins ATP6V1A, ATP6V1B2 and ATP6V1E1 were the subunits of vacuolar ATPase (v-ATPase). ATP6V1A was shown to be a catalytic subunit of the enzyme that hydrolyzed ATP whereas the other subunits played a role in interaction with cytoskeletal proteins and aldolases (like overexpressed ENO1 that we already mentioned in connection with Piruzian’s paper), which were the major suppliers of ATP for v-ATPase. In the lysosomes, v-ATPases were needed to maintain the proper pH in their lumen. Four other identified lysosomal proteins were cathepsins (CTSD and CTSG) and endogenous inhibitors of cathepsins (CSTA and CSTB). The observed changes in their expression suggested us a balance shift between the lysosomal hydrolases due to a decreased expression of CTSG, inhibition of CTSB, CTSH and CTSL by CSTA and CSTB and increased expression of CTSD. The suggested misbalance of lysosomal proteins according to the others [26] could be linked to an activation of mTORC pathway (see below) and be partially responsible for incomplete degradation of the nucleus and other intracellular organelles in lesional epidermal keratinocytes.

Unfortunately, some results of the PO analysis could be easy misinterpreted. For instance, we reported above that melanosome were strongly enriched for DEPs exhibited increased expression in lesional skin. However, a closer look on these proteins revealed that they were neither restricted to melanosome nor involved in the metabolism of melanin. Moreover, membrane-bound vesicles that we discussed before were also enriched for these DEPs.

In contrast to normally looking skin (Table S4), the extracellular components (exosomes, microparticles extracellular matrix etc.) were not enriched in lesional skin. However, similar terms, such as extracellular region (P = 1.12 x 10^−16^), extracellular matrix (P = 3.16 x 10^−10^) and extracellular space (P = 5.33 x 10^−8^) were enriched when we analyzed DEPs that exhibited decreased expression separately (Table S6b). This observation suggested us that two different scenarios could be implemented simultaneously to prepare the epidermis for remodeling: one – to modulate the developing inflammatory response in the extracellular space and another – to accelerate the protein metabolism in the cells affected by the disease.

The analysis involving molecular function terms (Table S8) showed the strongest enrichment for structural constituent of ribosome (P = 9.37 x 10^−40^), structural molecule activity (P = 9.29 x 10^−33^) and RNA binding (P = 5.4 x 10^−23^). This result did not surprise us, since proteins linked to translation, such as ribosome proteins were increased in lesional skin as we already showed above (Fig. 2). We also found enrichment for unfolded protein binding (P = 4.25 x 10^−7^), ATPase (P = 7.85 x 10^−5^) and RNA helicase activities. The DOPs that exhibited an increased expression in lesional skin were enriched in the same terms (Table S8a). The only new molecular function that we identified for this set of proteins was GTPase activity (P = 0.039). In contrast, the DEPs that exhibited decreased expression in lesional skin (Table S8b) were mainly enriched for glycosaminoglycan binding (P = 3.82 x 10^−7^), phosphatidylcholine-sterol O-acyltransferase activator activity (P = 1.15 x 10^−5^) and extracellular matrix structural constituent binding (P = 1.74 x 10^−5^) highlighting the link between decreased expression of proteins involved in binding to proteoglycans in lesional skin and the progression of the inflammatory response.

The DEPs that were involved in unfolded protein binding were represented by four groups of chaperones, TRiC, HSP70, HSP90 proteins and nuclear chaperones. In normally looking skin (Fig. 5), the expression levels of HSP70s (HSPA4, HSPA5 and HSPA8), HSP90s (HSP90AA1 and HSP90AB1), nuclear chaperones (NAP1L4, NPM1 and RUVBL2), co-chaperones (CDC37 and DNAJB1) and mitochondrial chaperone HSPD1 were decreased. In contrast, their expression levels were elevated in lesional skin regardless their location and role in the cell. The latter could be done to support accelerated mRNA translation and minimize losses due to protein misfolding in the cells affected by the disease.

**Figure 5.**
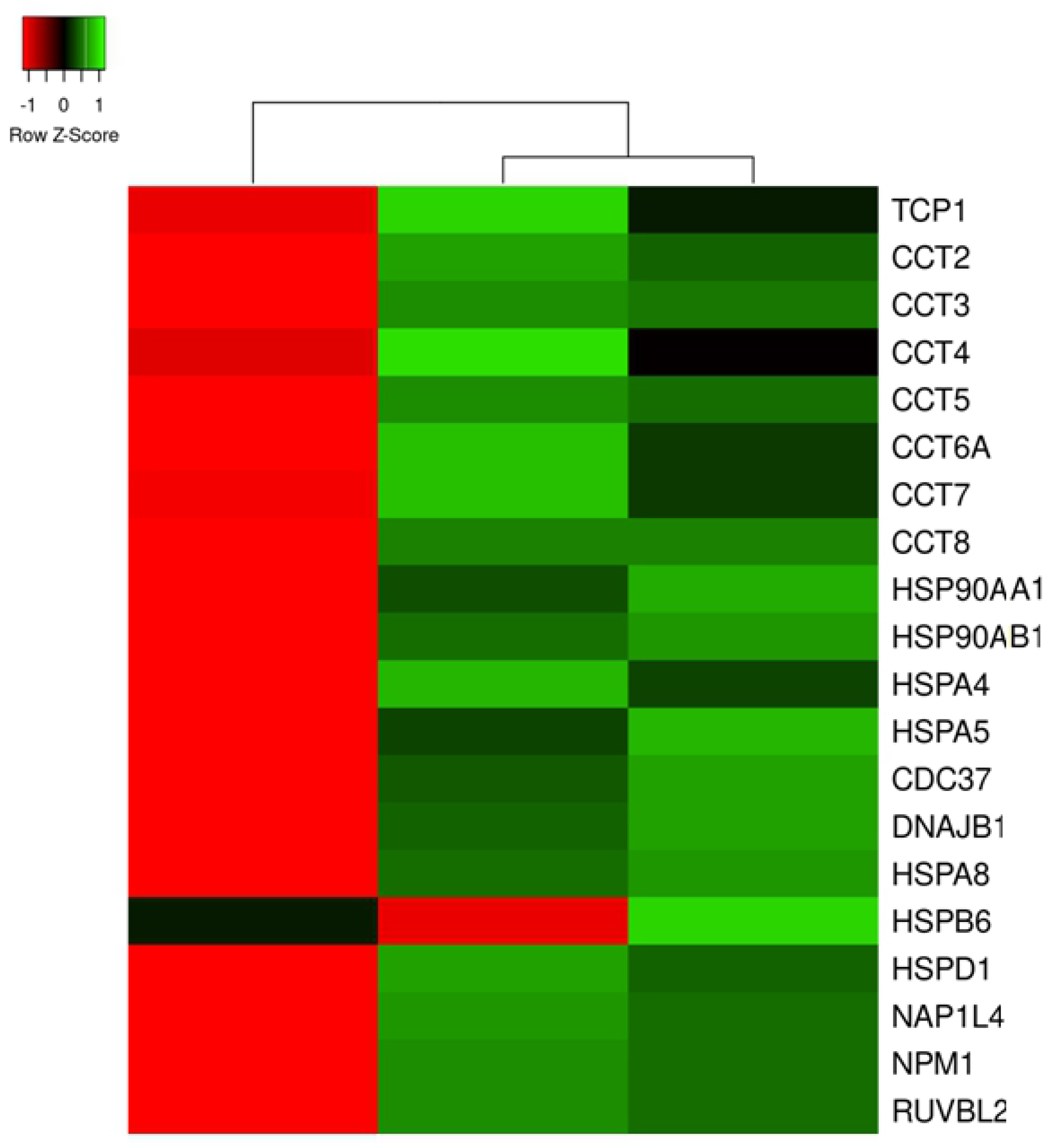
Heat map for the expression levels of the DEPs that function as chaperones in the cell. From left to right column – the results of comparative analysis for normally looking skin of the patients vs. skin of the healthy volunteers, lesional vs. normally looking skin of the patients, and lesional skin of the patients vs. skin of the healthy volunteers. Red color indicates low expression levels whereas green color indicates high expression levels. The DEPs uniprot ID, official gene symbols of the encoding genes, fold changes and Benjamini corrected P-values are shown in Supplementary Table S2.

Particularly, we discovered that all TRiC subunits (TCP1 and CCT1-8) exhibited the elevated expression levels. Considering the primary TRiC function in the cell to fold misfolded cytoskeletal proteins, such as actin and tubulin as well as the components of mTORC complexes [27] we proposed that us that lesional skin experienced an increased demand for protein folding. Earlier Swindell et al. [9] suggested that mTOR, the key component of mTORC, was governing the hypertranslation in psoriasis. In this regard, their hypothesis is supported by our findings that there was more TRiC complexes in lesional skin In turn, mTORC complexes were needed to organize the microfilaments and other cytoskeletal proteins in the cells and mTORCs activity supposed to be higher due to the ability of TRiC to activate mTORCs.

Notably, the expression profile of small chaperone HSPB6 was different from the others. Particularly, the expression level of HSPB6 was decreased in lesional skin and increased – in normally looking skin of the patients (Fig. 5). To date, it still not much is known of the role that HSPB6 may play in the lesional cell. Previously, it was reported that HSPB6, a cytoplasmic co-chaperone of HSP70, was activated by oxidative stress [28] and had the capability to sequester cytochrome C, which was leaked by the mitochondria to the cytoplasm to cause the apoptosis [29]. In our opinion, its downregulation in lesional skin could be driven by the necessity of shifting the balance between apoptosis and autophagy of intracellular organelles and toward autophagy and increasing the cell survival rate in the epidermis.

The rapidly proliferating psoriatic epidermis is on a high demand of energy. In turn, the cellular ATPases and ATP synthase are primary consumers and suppliers of energy in the cell. In this regard, we would like to mention a large group of DEPs, constituents of these enzymes, namely ATP1A1 and ATP1B3 (α_1_β_3_ Na^+^/K^+^ ATPase), ATP2A2 (the Ca2^+^-ATPase SERCA2), ATP6V1A, ATP6V1B2 and ATP6V1E1 (the subunits of vacuolar ATPases) and ATP5F1A/α-F_1_, ATP5F1B/β-F_1_, ATP5F1C/γ-F_1_, ATP5J/F6-F_0_, ATP5MF/f-F_0_, ATP5MG/g-F_0_ and ATP5PB/b-F_0_ (the subunits of ATP-synthase). All named proteins exhibited higher expression in lesional skin and lower expression in normally looking skin of the patients, compared to control (Fig. 6a) suggesting higher rates for both – ATP consumption and production. Particularly, it could be needed to achieve a faster turnover of epidermal keratinocytes during their shortened life cycle. Moreover, this finding is in a good agreement with our hypothesis that changes in psoriatic proteome were aimed to maintain a higher metabolic rate in lesional skin.

**Figure 6.**
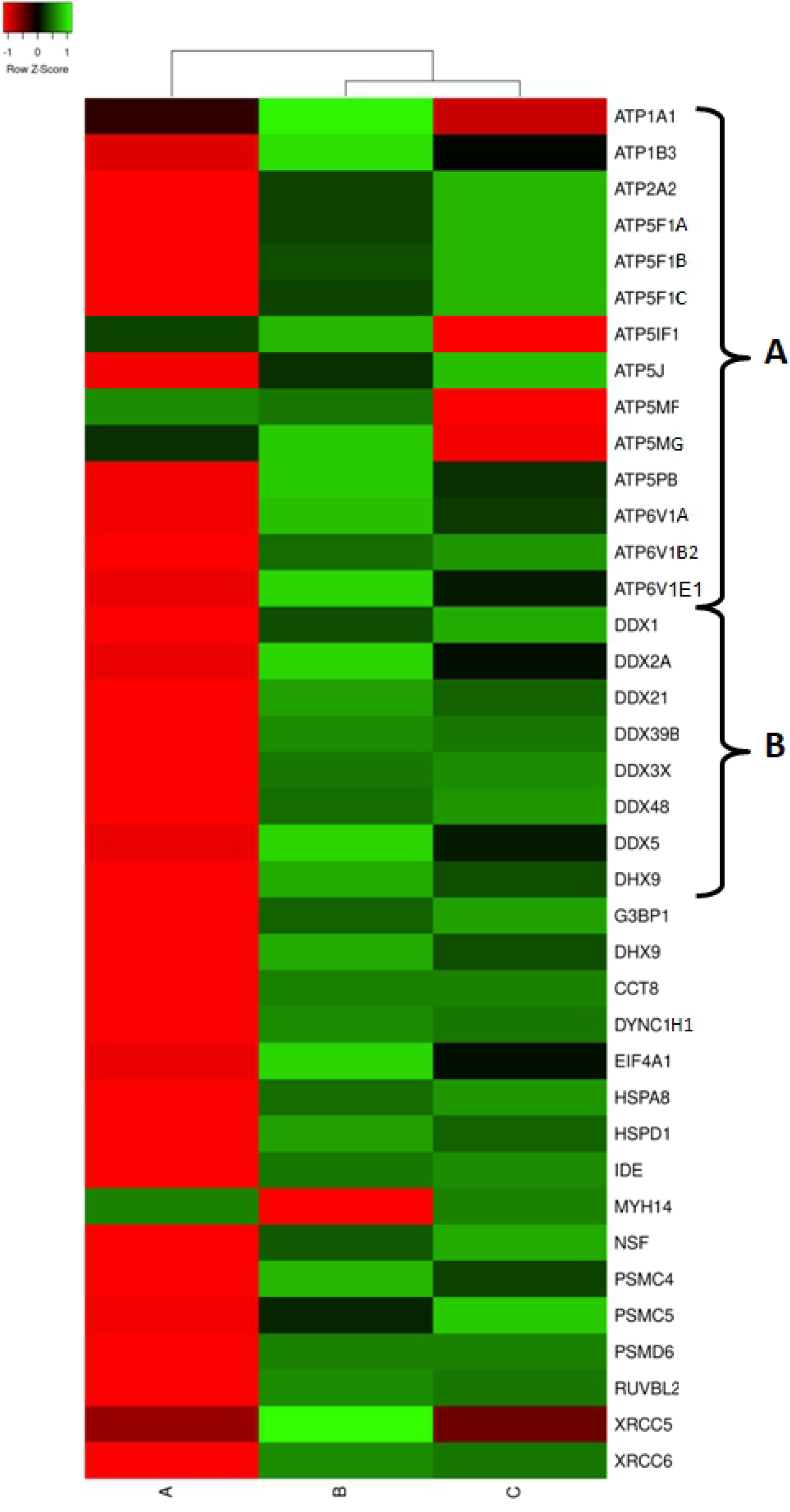
Heat map for the expression levels of DEPs capable of ATP binding. A – proteins constituting ATP synthase and ATPases. B. – RNA helicases. From left to right column – the results of comparative analysis for normally looking skin of the patients vs. skin of the healthy volunteers, lesional vs. normally looking skin of the patients, and lesional skin of the patients vs. skin of the healthy volunteers. Red color indicates low expression levels whereas green color indicates high expression levels. The DEPs uniprot ID, official gene symbols of the encoding genes, fold changes and Benjamini corrected P-values are shown in Supplementary Table S2.

In addition, we discovered a group of RNA helicases that were also upregulated in lesional skin and downregulated in normally looking skin of our patients (Fig. 6b). Due to the term “RNA helicases” seems to be very restrictive we have to mention that RNA helicases are associated with many aspects of RNA metabolism and functions ranging from transcription and translation, to processing and decay [30]. Since RNA metabolism as we showed above was deeply affected in lesional skin (Fig. 2), it was not surprising that RNA helicases exhibited a similar expression pattern. Expectedly, the differentially expressed RNA helicases identified in our study were deeply involved in any part of RNA metabolism that we named above. DDX1 was found to regulate the translation of insulin mRNA [31]. DDX3 was shown to regulate mRNA transcription [32, 33], splicing [34], participate in translocation of spliced mRNA to the cytoplasm [35, 36] and influence the translation [36-38]. Moreover, DDX3 was implicated in Drosha-mediated processing of pri-miRNAs [39] and regulation of cell cycle [32, 40, 41]. DDX5 was discovered to unwind miRNA precursor duplex to facilitate the loading of guide RNA on to miRISCs [42]. DDX6 was previously demonstrated to participate in the repression of translation, mRNA storage and decay [43-45]. DHX9 was shown to influence DNA replication, mRNA transcription, processing, exporting and translation [46]. Moreover, it was reported that DHX9 was capable of binding and resolving RNA G4 structures [47]. DDX39B was shown to have an active role in exporting the spliced mRNA from the nucleus to the cytoplasm [48]. Finally, DDX48/EIF4A3 was found to participate in pre-RNA splicing [49]. In addition, DDX48 was also demonstrated to unwind the 5’ untranslated region in the cytoplasm [50]. Similarly to the other groups of DEPs that we reviewed above the differentional expression of the named RNA helicases was needed to enforce rapid changes in the proteome necessary to maintain a higher metabolic rate in lesional skin.

In conclusion, performing LC-MS/MS analysis of skin samples obtained from psoriasis patients and healthy volunteers, we identified 520 DEPs that exhibited higher and lower expression levels in lesional and normally looking skin of psoriasis patients compared to the skin of the healthy volunteers. The following PO analysis revealed an activation of opposite metabolic processes, such as proteins biosynthesis and protein degradation. Analyzing the proteins differentially expressed in lesional skin, we discovered 54 ribosome proteins, 10 aminoacyl-tRNA synthetases, 10 RNA helicases and 21 protein factors controlling initiation elongation and termination of translation those expression was increased. On the other hand, we detected ∼20 molecular chaperones and about the same number proteasome subunits those expression levels were also elevated. We also found an increased expression of the proteins that constitute ATP-synthase and ATPases, the major suppliers and consumers of ATP in the cell. In addition, we reported of an increased expression of proteins that maintain the normal work of protein degrading cell organelles – endosomes and lysosomes. Collectively, our results explain the molecular basis of a higher metabolic rate in psoriatic skin lesions and provide new insights to the metabolic processes affected by the disease.

